# cyAbrB transcriptional regulators as safety devices to inhibit heterocyst differentiation in *Anabaena*

**DOI:** 10.1101/539908

**Authors:** Akiyoshi Higo, Eri Nishiyama, Kota Nakamura, Yukako Hihara, Shigeki Ehira

## Abstract

Cyanobacteria are monophyletic organisms that perform oxygenic photosynthesis. While they exhibit great diversity, they have a common set of genes. However, the essentiality of them for viability has hampered the elucidation of their functions. One example of the genes is *cyabrB1* encoding a transcriptional regulator. In the present study, we investigated the function of *cyabrB1* in heterocyst-forming cyanobacterium *Anabaena* sp. PCC 7120 through CRISPR interference, a method we recently utilized for the photosynthetic production of a useful chemical in the strain. Conditional knockdown of *cyabrB1* in the presence of nitrate resulted in formation of heterocysts. Two genes, *hetP* and *hepA*, which are required for heterocyst formation, were up-regulated by *cyabrB1* knockdown in the presence of combined nitrogen sources. The genes are known to be induced by HetR, a master regulator of heterocyst formation. *hetR* was not induced by *cyabrB1* knockdown. *hetP* and *hepA* were repressed by direct binding of cyAbrB1 to their promoter regions in a HetR-independent manner. In addition, the over-expression of *cyabrB1* abolished heterocyst formation upon nitrogen depletion. Also, knockout of *cyabrB2*, a paralogue gene of *cyabrB1,* in addition to *cyabrB1* knockdown, enhanced heterocyst formation in the presence of nitrate, suggesting functional redundancy of cyAbrB proteins. We propose that a balance between amounts of HetR and cyAbrB1 is a key factor influencing heterocyst differentiation during nitrogen step-down. cyAbrB proteins are essential safety devices inhibiting heterocyst differentiation.

**Importance:** Spore formation in *Bacillus subtilis* and *Streptomyces* represents non-terminal differentiation and has been extensively studied as models of prokaryotic cell differentiation. In the two organisms, many cells differentiate simultaneously, and the differentiation is governed by a network in which one regulator stands at the top. Differentiation of heterocysts in *Anabaena* sp. PCC 7120 has also been extensively studied. The differentiation is unique because it is terminal and only 5-10% vegetative cells differentiate into heterocysts. In the present study, we identified cyAbrB1 as a repressor of two genes that are essential for heterocyst formation, *hetP and hepA*, independent of HetR, which is a master activator for heterocyst differentiation. The finding is reasonable for unique cell differentiation of *Anabaena* because cyAbrB1 could suppress heterocyst differentiation tightly in vegetative cells, while only cells in which HetR is over-expressed could differentiate into heterocysts.

## Introduction

Cyanobacteria are ancient and monophyletic prokaryotes, which are characterized by a capacity to perform oxygenic photosynthesis. They are found in diverse habitats, including fresh and marine water, hot springs, frozen lakes, soil, and deserts (1). Specific responses to environmental changes enable them to adapt to their habitats (2). In addition, they exhibit a great diversity of morphology and cell arrangements. Moreover, some cyanobacteria can differentiate into specific cell types in response to environmental stimuli, which is one type of stress response. The most studied differentiated cell type in cyanobacteria is the heterocyst. At semi-regular intervals, some filamentous cyanobacteria can differentiate into larger and round cells called heterocysts, which are cells specialized for nitrogen fixation, which enables heterocystous cyanobacteria to inhabit nitrogen-poor environments.

*Anabaena* sp. PCC 7120 (*Anabaena*) has been extensively studied as a model for heterocyst differentiation (3,4). Upon the depletion of combined nitrogen, 5-10% of vegetative cells that perform oxygenic photosynthesis differentiate into heterocysts. A transcriptional regulator, NtcA, widely conserved in cyanobacteria, perceives nitrogen deficiency as an increase of a metabolite 2-oxoglutarate (5). Subsequently, NtcA indirectly induces HetR, a master regulator of heterocyst differentiation (6). Accumulation of HetR spatially initiates specific developmental program and enables patterned heterocyst formation (7–9). During differentiation, deposition of exopolysaccharide and glycolipid layers results in morphological changes in the cells. In addition, cellular metabolism is dynamically altered by the inactivation of oxygenic photosystem II and enhancement of respiration (3,4). Such changes enable heterocysts to protect oxygen-labile nitrogenase from oxygen.

Despite their great diversity, cyanobacteria have a core set of genes that are conserved across the phylum (10, 11). Many of the conserved genes have been found to be associated with core biological process such as DNA replication, transcription, translation, photosynthesis, the Calvin cycle, and various metabolic pathways (11). Therefore, many of their functions can be predicted. However, the functions of some of the conserved genes are yet to be elucidated. Although the study of such genes could offer novel insights into cyanobacterial biology, essentiality of the core genes (11, 12) has hampered such investigations. An example of the genes is *cyabr1* (13).

*cyabrB* encoding a transcriptional regulator is conserved among cyanobacteria (13). The DNA-binding domain of cyAbrB located at the C-terminus belongs to AbrB-like family (Pfam14250), which is unique to cyanobacteria and is not conserved in other organisms including melainabacteria, a non-photosynthetic sister phylum to the cyanobacteria. Each cyanobacterium has two copies of *cyabrB* genes, *cyabrB1* and *cyabrB2* (also known as *calA* and *calB* in *Anabaena*, respectively). Because some attempts to disrupt *cyabrB1* in *Anabaena* and unicellular cyanobacterium *Synechocystis* sp. PCC 6803 (*Synechocystis*) have failed so far (13–16), the gene should be essential for cyanobacteria. Conversely, gene disruption of *cyabrB2* is possible in *Synechocystis*. Studies on *cyabrB2* mutants have revealed that *cyAbrB2* is involved in acclimation to changes in carbon and nitrogen availability (13, 17–19). While the studies demonstrated that *cyabrB2* had specific functions, some evidence suggested a functional overlap between cyAbrB1 and cyAbrB2 (20).

Recent studies towards photosynthetic production of useful chemicals have rapidly developed tools for artificial gene regulation systems for cyanobacteria (21–23). Among them, a gene knockdown technology, CRISPR interference (CRISPRi), has attracted much attention because the system exhibited repression over a wide dynamic range in an inducer concentration-dependent manner (24–28). Target genes can be repressed by the formation of a complex consisting of a nuclease-deficient Cas9 (dCas9), a single guide RNA (sgRNA), which corresponds to the target DNA sequence, and target DNA (29). While CRISPRi has been successfully applied to enhance the production of desired products in cyanobacteria (24–26, 30), its basic scientific applications are still awaited.

A question of whether a transcriptional regulator cyAbrB1 conserved in cyanobacteria regulates core genes or specific genes in *Anabaena* motivated us to study the function of *cyabrB1*. In the present study, we created *Anabaena* strains in which *cyabrB1* is conditionally knocked down through CRISPRi technology. cyAbrB1 amounts were significantly repressed in any conditions tested and in any genetic background tested when the CRISPRi system was induced. Repression of *cyabrB1* resulted in formation of heterocysts even in the presence of nitrate. Not *hetR*, but two direct target genes of HetR, *hetP* and *hepA,* which are required for heterocyst development (4), were induced by *cyabrB1* knockdown in the presence of combined nitrogen in a HetR-independent manner. Over-expression of *cyabrB1* abolished heterocyst formation under nitrogen-depleted conditions. Therefore, we concluded that cyAbrB1 is essential for the suppression of heterocyst differentiation and propose a model that cyAbrB1 offers HetR an appropriate threshold for the induction of heterocyst development.

## Results

### Heterocyst formation by cyabrB1 knockdown in the presence of nitrate

A conditional *cyabrB1* knockdown strain C104 was constructed by integrating a plasmid containing *P*_*petE*_-*tetR*, *P*_*L03*_-*dcas9*, and *P*_*J23119*_-sgRNA targeting *cyabrB1* (Figure S1) to the neutral site *cyaA* (31). In the strain, an inducer anhydrotetracycline (aTc) derepresses *L03* promoter by binding to TetR, and *dcas9* is induced. sgRNA is constitutively expressed. Therefore, the addition of aTc switches on repression by CRSIPRi. A negative control strain C100 without sgRNA was also constructed.

Strain C104 was grown in nitrate-containing medium and bubbled with air containing 1% (v/v) CO_2_. When the expression of *cyabrB1* was repressed for 48 h, 0.6% of heterocysts were formed (Fig. S2A). In contrast, when *cyabrB1* was not repressed, little heterocysts (<0.1%) were formed. To facilitate clearer observation of the phenotype, C104 was cultured in nitrate-containing medium and bubbled with air containing 5% (v/v) CO_2_, in which carbon is excess to nitrogen in the cells. In this condition, C104 and C100 formed 0.5 and 0.4% of the heterocysts, respectively, in the absence of the inducer. In the presence of the inducer, C104 formed 2.9% of the heterocysts but C100 did not form any heterocysts (Fig. 1A) (<0.1%). Knockdown of cyAbrB1 was confirmed by western-blotting analysis (Fig. 1B and C). While an addition of aTc to control strain C100 did not repress cyAbrB1 expression, in the case of C104, addition of aTc considerably repressed cyAbrB1 expression (less than 10%), particularly at 48 and 72 h. Repression of *cyabrB1* did not lead to heterocyst formation when C104 was cultured in ammonium-containing medium (Fig. S2B).

**Fig. 1.**
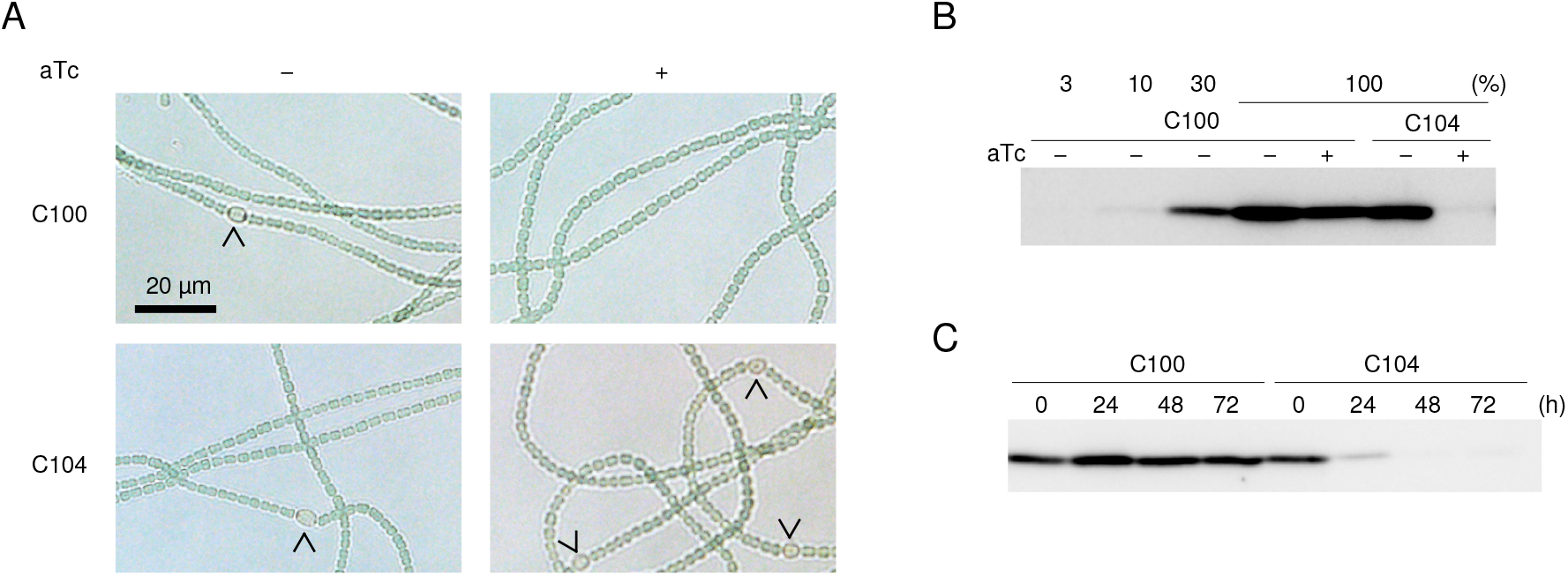
Promotion of heterocyst formation by knockdown of *cyabrB1* in the presence of nitrate. Control strain lacking sgRNA (C100) and *cyabrB1* knockdown strain (C104) were cultured in nitrate-containing medium in the absence or presence of the inducer aTc (50 ng/ml). Cultures were bubbled with air containing 5% (v/v) CO_2_. (A) Formation of heterocysts by *cyabrB1* knockdown in the presence of nitrate. Images were photographed after 48 h of cultivation. Arrowheads indicate heterocysts. (B) Confirmation of cyAbrB1 knockdown. After 48 h of cultivation, total protein was extracted and western blotting using anti-cyAbrB1 antibody was performed. Different amounts of total proteins from C100 cultured in the absence of aTc were loaded to show the linearity of the results. (C) Time course analysis of cyAbrB1 knockdown. Each strain was cultivated for the indicated time in the presence of 50 ng/ml aTc. Subsequently, total protein was extracted and western blotting using anti-cyAbrB1 antibody was performed.

To rule out the possibility that the observed phenotype is due to off-target effects of CRISPRi, we constructed C105 and C106 that retain sgRNA targeting different sites of *cyabrB1*. Formation of heterocyst was promoted in both C105 and C106 by *cyabrB1* repression similarly in C104, when cells were cultured in nitrate-containing medium and bubbled with air containing 5% (v/v) CO_2_ (Fig. S3). The results suggest that cyAbrB1 is required for the suppression of heterocyst formation in the presence of nitrate.

### Repression of *hetP* and *hepA* by cyAbrB1 in the presence of nitrogen sources

To elucidate the cause of heterocyst formation in the presence of nitrate, RNA was extracted from cells of C100 and C104 cultured for 48 h in nitrate-containing medium with 5% CO_2_ in the absence or presence of the inducer, and an RT-qPCR analysis was performed. The expression of four genes that are induced at early stages of heterocyst development, including *hetR, hetP, hetZ*, and *hepA*, was investigated (Fig. 2A). *hetR* encodes a master regulator of heterocyst differentiation and its over-expression causes heterocyst formation in the presence of nitrogen sources (9). Both *hetP* and *hetZ* are directly up-regulated by HetR, and ectopic expression of either gene leads to heterocyst formation in the presence of nitrogen sources (4, 32–36). *hepA* encodes a component of an ABC transporter required for the construction of the heterocyst exopolysaccharide layer (37), which is the first step during morphological differentiation (4). HetR also directly induces expression of *hepA* (4, 38). While the expression of *hetR* and *hetZ* did not change significantly, the expression of *hetP* and *hepA* was significantly induced when *cyabrB1* was repressed in C104. Similar results were observed in C105 and C106 (Fig. 2A). We confirmed that *cyabrB1* transcript and cyAbrB1 protein in C104, C105, and C106 were repressed in the presence of the inducer (Fig. S4A and B, respectively).

**Fig. 2.**
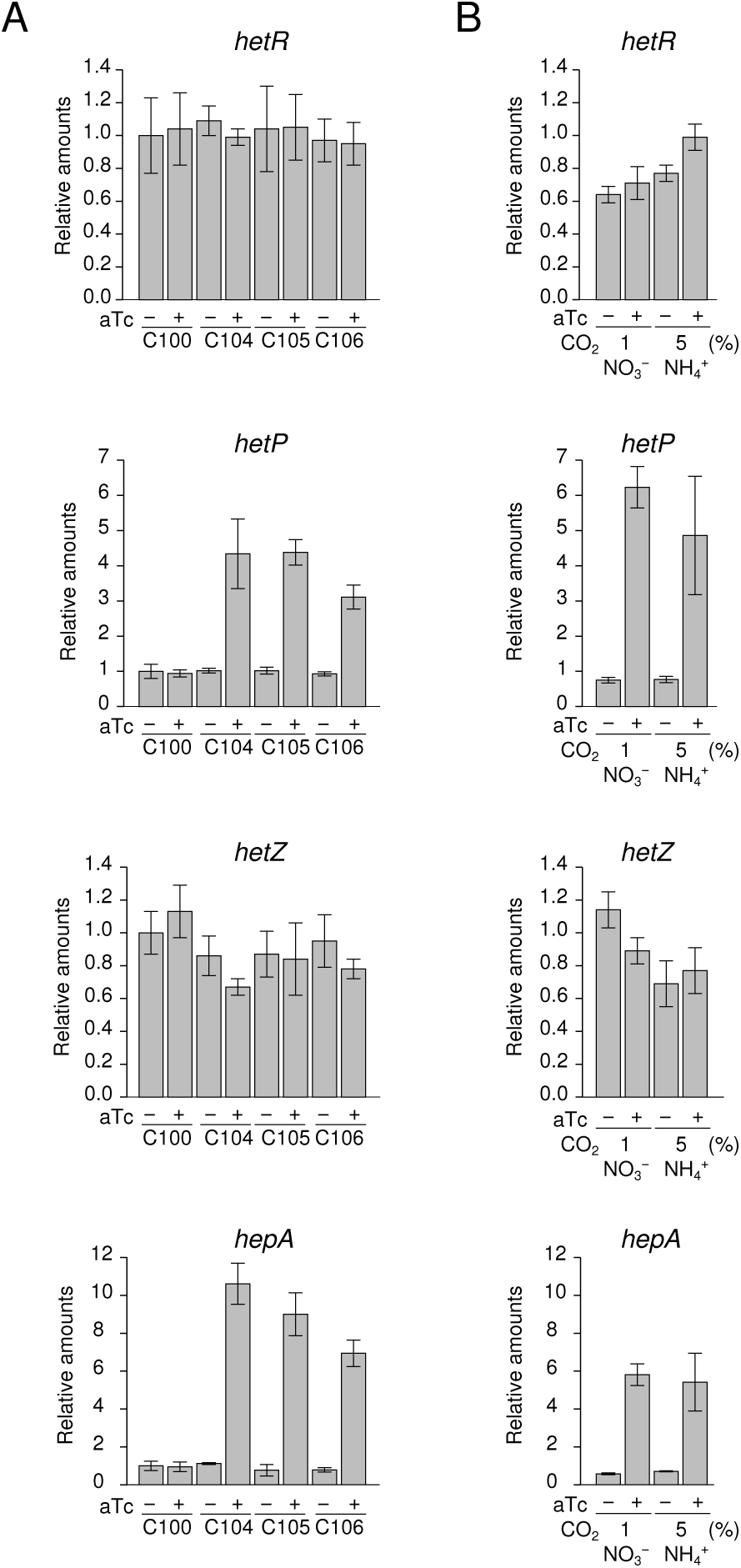
Up-regulation of some genes related to heterocyst differentiation following knockdown of *cyabrB1*. After 48 h of cultivation in the absence or presence of 50 ng/ml aTc, RNA was extracted and RT-qPCR was performed. *rnpB* was used for normalization. Data represent mean ± SD (n = 3 from independent culture). Amounts of each gene relative to that in C100 grown in nitrate-containing medium bubbled with air containing 5% (v/v) CO_2_ in the absence of the inducer are shown. (A) Each strain was grown in nitrate-containing medium and bubbled with air containing 5% (v/v) CO_2_. (B) Strain C104 was grown in nitrate-containing medium bubbled with air containing 1% (v/v) CO_2_, or in ammonium-containing medium bubbled with air containing 5% (v/v) CO_2_.

Subsequently, we extracted RNA from C104 cultured in the presence of nitrate with 1% CO_2_ bubbled or in the presence of ammonium with 5% CO_2_ bubbled (Fig. 2B). Although no heterocysts were formed in the latter condition, *hetP* and *hepA* were greatly induced when *cyabrB1* was repressed in both conditions, similarly in Fig 2A. We confirmed that *cyabrB1* transcript and cyAbrB1 protein were repressed in both conditions in the presence of the inducer (Fig. S4C and D, respectively). The results indicate that the up-regulation of *hetP* and *hepA* by *cyabrB1* knockdown induced heterocyst formation rather than *hetP* and *hepA* were induced following the initiation of heterocyst development, and that heterocyst formation was suppressed by an unknown mechanism in the presence of ammonium.

Expression of *nifH* encoding a subunit of nitrogenase was quantified to determine whether the heterocysts formed by *cyabrB1* repression were functional. Repression of *cyabrB1* in C104 in the presence of nitrate bubbled with 5% CO_2_ induced *nifH*, but not under bubbling with 1% CO_2_ (Fig. S5), suggesting that maturation of heterocysts depends on C-N balance inside the cells and does not directly depend on cyAbrB1. Therefore, we concluded that cyAbrB1 is essential for repression of *hetP* and *hepA* in the presence of nitrogen sources.

### Induction of *hetP* and *hepA* by cyAbrB1 knockdown independently of HetR

To clarify whether the up-regulation of *hetP* and *hepA* is independent of or dependent on HetR, we constructed a C104h strain in which *cyabrB1* could be knocked down using the CRISPRi system in a *hetR* mutant strain (39) (Fig. S1). RNA was extracted from C104h cultured in the presence of nitrate bubbled with 5% CO_2_ in the absence or presence of aTc. RT-qPCR analysis revealed that *hetP* and *hepA* were up-regulated by *cyabrB1* repression in the *hetR*-deficient background (Fig. 3A), indicating that cyAbrB1 regulates the expression of the two genes independently of HetR. We confirmed that *cyabrB1* transcript and cyAbrB1 protein in C104h were repressed similarly in C104 in the presence of the inducer (Fig. 3A and B).

**Fig. 3.**
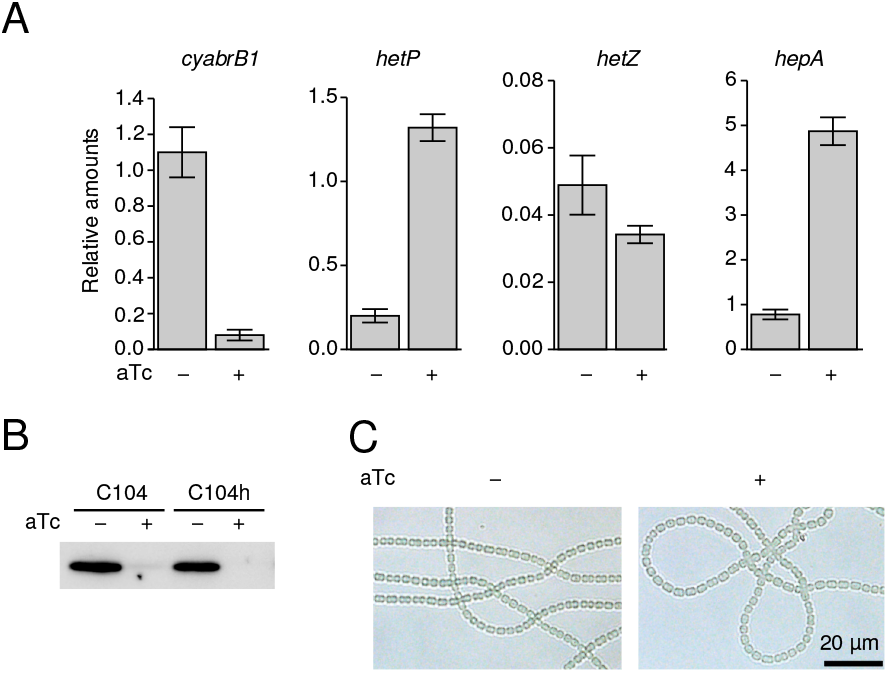
Knockdown of *cyabrB1* in the *hetR* mutant background. Cells were cultured in nitrate-containing medium bubbled with air containing 5% (v/v) CO_2_ in the absence or presence of aTc for 48 h. (A) Effect of *hetR* inactivation on expression of *cyabrB1* and genes related to heterocyst formation. RNA was extracted and RT-qPCR was performed. *rnpB* was used for normalization. Data represent mean ± SD (n = 3 from independent culture). Amounts of each gene relative to that in C100 cultured in nitrate-containing medium bubbled with air containing 5% (v/v) CO_2_ in the absence of the inducer are shown. (B) Confirmation of cyAbrB1 knockdown. Total protein was extracted and western blotting using anti-cyAbrB1 antibody was performed. (C) No heterocyst formation by *cyabrB1* knockdown in *hetR* deletion background. Cells of C104h were micro-photographed.

Heterocysts were not formed following the repression of *cyabrB1* in C104h (Fig. 3C). Although the expression of *hetP* in C104h was induced by *cyabrB1* in the presence aTc compared to in the absence of the inducer, the expression levels of *hetP* in the presence of the inducer in C104h were low compared to those in C104 (Figs. 2A and 3A), possibly due to the effect of *hetR* disruption. The result could explain why heterocysts were not formed even when *hetP* and *hepA* were induced in C104h (Fig. 3C).

### Specificity and redundancy of cyAbrB proteins

Subsequently, we constructed a *cyabrB2* knockout mutant DR2080 and a *cyabrB1* knockdown/*cyabrB2* knockout mutant C104B2 (Fig. S1) to examine specificity and redundancy of cyAbrB1 and cyAbrB2. In C104B2, *cyabrB1* knockdown caused 4.5% heterocyst formation in nitrate-containing medium bubbled with 1% CO_2_ (Fig. 4A). In contrast, *cyabrB1* knockdown mutant C104 and *cyabrB2* knockout mutant DR2080 produced only 0.6 and 0% heterocyst under similar conditions, respectively (Fig. S2A and Fig. 4A). An RT-qPCR analysis revealed that *hetP* and *hepA* were similarly induced by *cyabrB1* knockdown in C104 and C104B2 (Compare Figs. 2B and 4B). The expression of *hetR* and *hetZ* were not induced by *cyabrB1* knockdown in C104B2, similarly to in C104 (Fig. 4B). Deletion of *cyabrB2* hardly influenced expression of *hetR, hetP, hetZ*, and *hepA*. Repression of *cyabrB1* transcript and cyAbrB1 protein in C104B2 was confirmed (Fig. S6A and B). C104B2 did not form heterocysts in ammonium-containing medium (Fig. S6C). Comparison of the results with *cyabrB1* knockdown, *cyabrB2* knockout, and *cyabrB1* knockdown/*cyabrB2* knockout mutants revealed that cyAbrB1 but not cyAbrB2, specifically regulates the expression of *hetP* and *hepA*. However, with regard to heterocyst formation in the presence of nitrate, cyAbrB1 and cyAbrB2 could be redundant since the double mutant produced more heterocysts compared to single mutants.

**Fig. 4.**
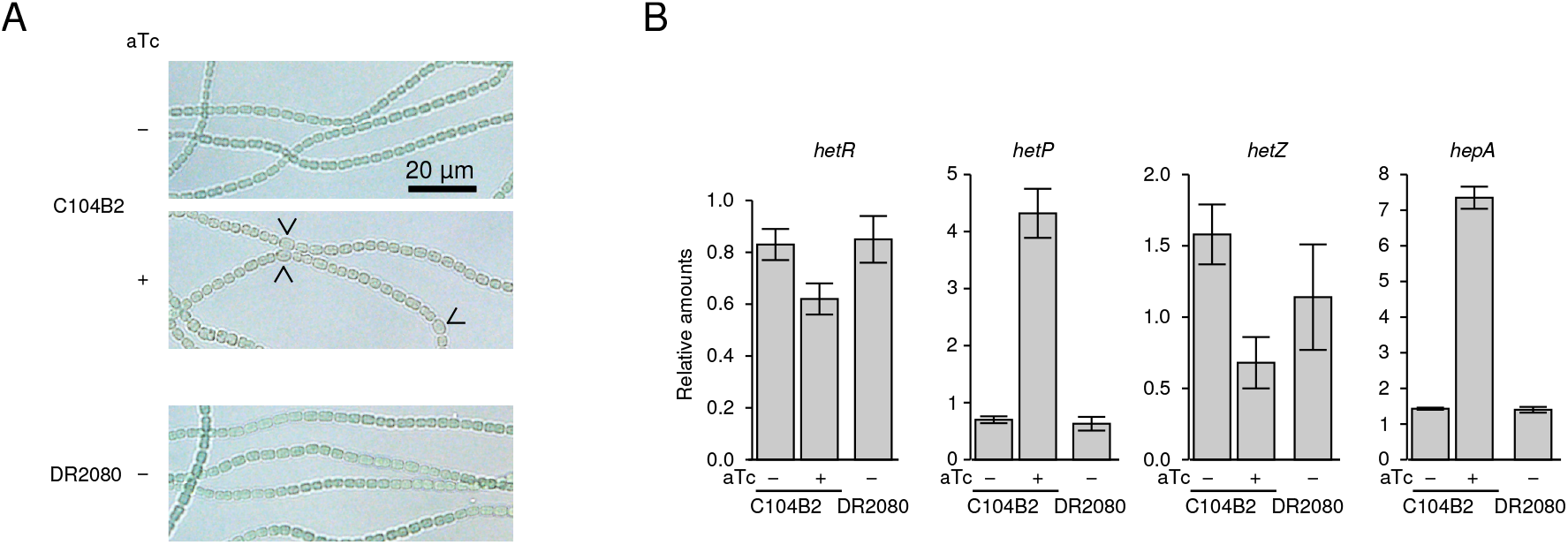
Specific and redundant functions of *cyabrB* genes. Cells were cultured in nitrate-containing medium bubbled with air containing 1% (v/v) CO_2_ in the absence or presence of aTc for 48 h. (A) Formation of heterocysts by *cyabrB1* knockdown and *cyabrB2* knock-out. Cells of C104B2 or DR2080 were micro-photographed. (B) RNA was extracted and RT-qPCR was performed. *rnpB* was used for normalization. Data represent the mean ± SD (n = 3 from independent culture). Amounts of each gene relative to that in C100 grown in nitrate-containing medium bubbled with air containing 5% (v/v) CO_2_ in the absence of the inducer are shown.

### Direct binding of cyAbrB1 to promoters of *hetP* and *hepA*

To test whether the expression of *hetP* and *hepA* was directly regulated by cyAbrB1, we expressed a recombinant His-cyAbrB1 in *Escherichia coli* and purified it. The purified His-cyAbrB1 protein had an apparent molecular weight of 18,000, which was largely consistent with the theoretical value (Fig. 5A). We performed a gel mobility shift assay using His-cyAbrB1 and Cy3-labeled DNA probe PhetP that includes *hetP* promoter region (Fig. 5B). His-cyAbrB1 retarded the mobility of the probe. Thereafter, we examined the specificity of the interaction using a competition assay. Addition of a 5 or 10-fold molar excess of non-labeled DNA probe PhetP and PhepA (*hepA* promoter region) eliminated the retardation, but that of cyabrB1RT (internal region of *cyabrB1*) did not (Fig. 5B). The results indicate that cyAbrB1 binds promoter regions of *hetP* and *hepA* and that the two genes are directly repressed by cyAbrB1.

**Fig. 5.**
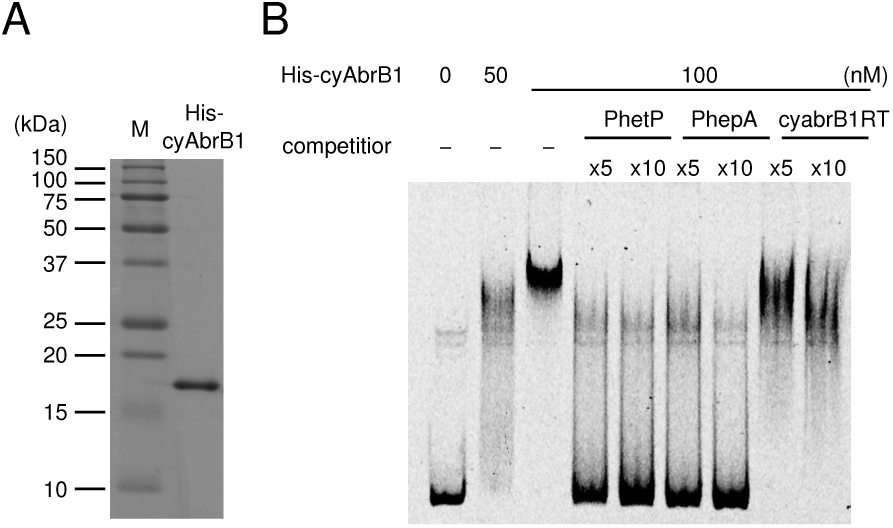
Direct binding of cyAbrB1 to promoters of *hetP* and *hepA*. (A) Purification of His-cyAbrB1. Purified His-cyAbrB1 was subjected to 15% SDS-PAGE. Lane M, protein molecular weight marker; lane His-cyAbrB1, purified His-cyAbrB1. (B) Electrophoresis mobility shift assay with His-cyAbrB1. cyAbrB1 was mixed with 3 nM DNA probe (promoter region of *hetP*). Non-labeled DNAs (PhetP, PhepA, or internal region of *cyabrB1* (cyabrB1RT)) were added.

### Inhibition of heterocyst formation by over-expression of *cyabrB1*

We investigated whether or how cyAbrB1 participates in heterocyst development in the absence of nitrogen sources. *cyabrB1* was knocked down upon removal of combined nitrogen sources in C104. After 24 or 48 h of nitrogen step-down, heterocysts were formed in the strain in the absence or presence of aTc (Fig. S7A and B). However, vegetative cell intervals were shorter in the presence of the inducer than in the absence of the inducer (Fig. S7C). Figure S7D shows that cyAbrB1 was repressed in the presence of the inducer in C104 at 48 h in the absence of nitrogen sources.

Then, we constructed a *cyabrB1* over-expression strain T121 (Fig. S1). In the strain, aTc induced the expression of *cyabrB1* driven by *P*_*L03*_. Strain C100, in which aTc induced the expression of *dcas9* but not *cyabrB1*, was used as the control strain. C100 or T121 was transferred from a nitrate-containing medium to a nitrogen-free medium, and was grown in the absence or presence of aTc. While C100 grew regardless of the absence or presence of aTc, T121 did not grow at all in the presence of the inducer (Fig. 6A). In contrast, the over-expression of *cyabrB1* only minimally inhibited the growth of T121 in the presence of nitrate, as previously demonstrated (40). Microscopic observations revealed that the addition of aTc to T121 abolished heterocyst formation 24 h after nitrogen depletion (Fig. 6B). Expression levels of *hetR, hetP*, and *hepA* were measured after depletion of nitrogen sources for 8 h, at which the genes were up-regulated in the wild type strain (41). While the expression of *hetP* was not repressed, the expression of *hepA* was repressed in T121 in the presence of aTc compared to in the absence of aTc, or in C100 in the absence or presence of aTc (Fig. 6B). Expression of *hetR* was also repressed in T121 following the addition of aTc, although it was minimal compared to the expression of *hepA* for an unknown reason. We confirmed that the addition of aTc led to the accumulation of *cyabrB1* transcripts and cyAbrB1 protein (Fig. 6B). The results suggested that over-production of cyAbrB1 inhibited the transcription of *hepA*. The reason why the expression of *hetP* was not inhibited is discussed below.

**Fig. 6.**
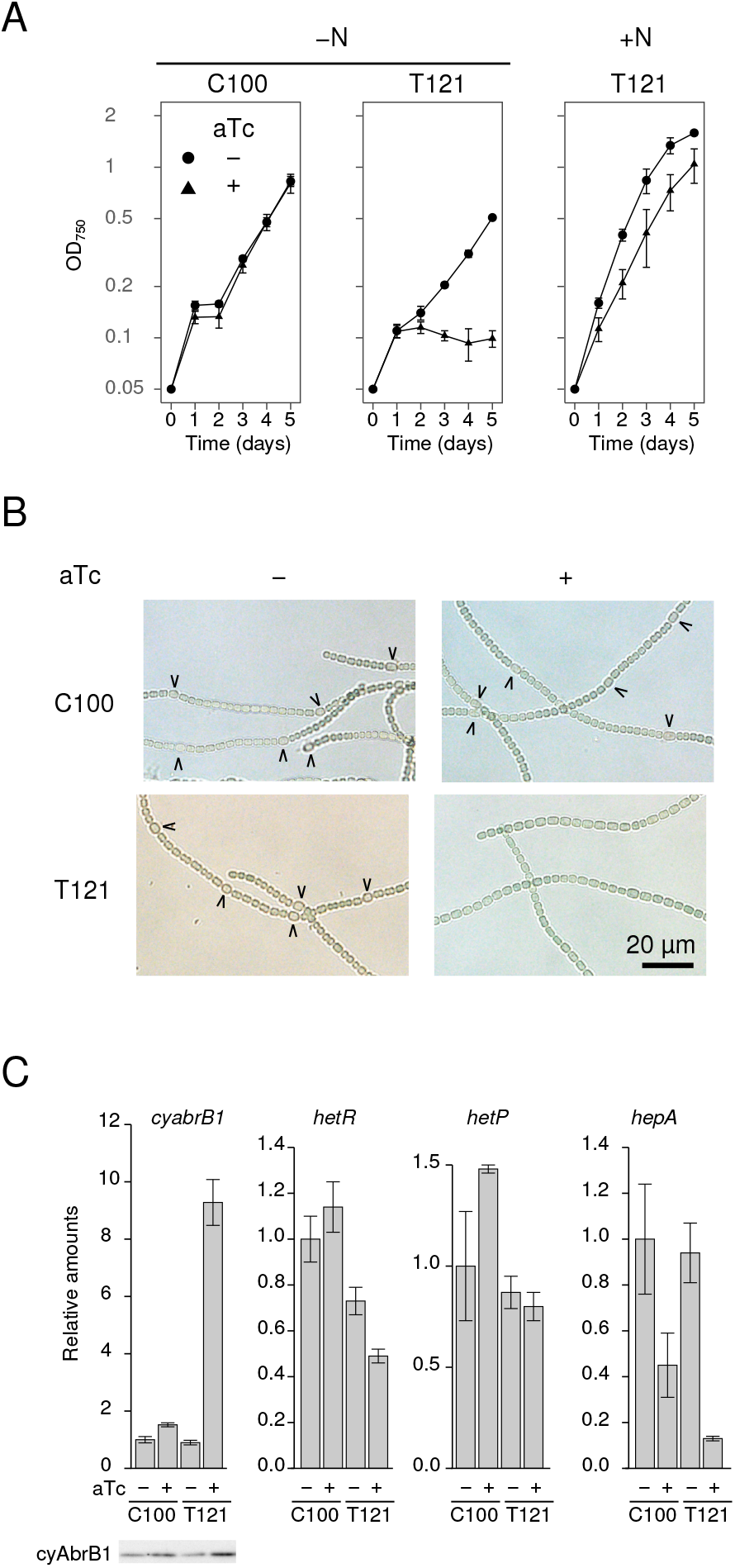
Effect of *cyabrB1* over-expression on heterocyst development. Cells of the negative control strain C100 and *cyabrB1* over-expression strain T121 were cultured in the absence or presence of 200 ng/ml aTc. (A) Impaired growth under nitrogen-depleted conditions by *cyabrB1* over-expression. Cells were cultured in nitrogen-free or nitrate-containing medium and OD_750_ was monitored. Data represent mean ± SD (n= 3 from independent culture). (B) Effect of *cyabrB1* over-expression on heterocyst formation. Cells of C100 and T121 cultured for 24 h in the absence or presence of the inducer were micro-photographed. (C) Effect of *cyabrB1* over-expression on expression of genes related to heterocyst formation. After 8 h of nitrogen depletion, RNA was extracted and RT-qPCR was performed. *rnpB* was used for normalization. Data represent mean ± SD (n = 3 from independent culture). Amounts of each gene relative to that in C100 grown in nitrate-free medium in the absence of the inducer are shown. Over-expression of cyAbrB1 was confirmed by western blotting.

We quantified the amounts of cyAbrB1 after nitrogen depletion in PCC 7120. Cells of PCC 7120 were washed by nitrogen-free medium and transferred to nitrate-containing or nitrogen-free medium, and were cultured for 8 h. Total proteins were extracted from cells before and after their cultivation. Western blotting using anti-cyAbrB1 antibody revealed that the depletion of nitrogen sources did not alter the amounts of cyAbrB1 (Fig. S8A and B). We also quantified the amounts of cyAbrB1 in mature heterocysts after nitrogen depletion for 24 h relative to that in whole filaments. The amounts of cyAbrB1 were marginally lower in heterocysts (Fig. S8C and D). The results suggested that high induction of HetR in proheterocysts (42) rather than decreasing cyAbrB1 amounts was a limiting step in the up-regulation of *hetP* and *hepA* during heterocyst development in the wild type strain following combined nitrogen step-down.

### Effect of *cyabrB1* repression or over-expression on the expression of *alr0947*

It has previously been shown that *cyabrB1* and *alr0947* constitute an operon (40). Therefore, we evaluated the polar effect of *cyabrB1* repression. While the expression of *cyabrB1* was highly repressed following the addition of aTc in C104, C105, and C106 in some conditions (Fig. S4A and C), the expression of *alr0947* was only slightly repressed (Fig. S9A and B). It was demonstrated that the over-expression of cyAbrB1 repressed the expression of *alr0947* since cyAbrB1 inhibits the transcription of the *cyabrB1-alr0947* operon (40). When *cyabrB1* was over-expressed, *alr0497* was repressed marginally (Fig. S9). While both repression and induction of *cyabrB1* resulted in a slight decrease in *alr0947* expression, repression induced heterocyst formation and the induction inhibited heterocyst formation, indicating that the observed phenotypes in the present study were caused by changes in expression levels of *cyabrB1* rather than those of *alr0947*.

## Discussion

In the present study, we revealed that cyAbrB1 is essential for the repression of *hetP* and *hepA* in the presence of nitrogen sources and that heterocysts are formed even in the presence of nitrate when *cyabrB1* is knocked out. Since the first description of *cyabrB1* in cyanobacteria (13), its essentiality has hampered investigation of the function of the gene. We overcome the challenge by applying CRISPRi, which is recently developed technology that has been employed to facilitate photosynthetic production of desired chemicals in some model cyanobacteria (24–26). Here, we demonstrated that the technology is also very useful for basic research. In all the conditions tested, and in any genetic backgrounds, CRISPRi facilitated the robust knock down of *cyabrB1*. The inhibition of heterocyst formation by the over-expression of *cyabrB1* in the absence of nitrogen sources has been overlooked so far. Agervald et al. (40) used the *nirA* promoter, which over-expresses only in the presence of nitrate while He and Xu (16) used the *petE* promoter, which might not be strong enough to hinder heterocyst formation. Consequently, the development of a variety of tools to regulate gene expression (23) is fundamental and could broaden the range of basic research and biotechnological application.

### Balance between cyAbrB1 and HetR determines heterocyst differentiation

Some evidence indicated that cyAbrB1 hinders heterocyst formation in the presence of nitrogen sources by repressing the expression of *hetP* and *hepA* through specific binding to promoter regions of the genes, which are direct targets of HetR. The possibility that the up-regulation of the genes by *cyabrB1* knockdown is due to an increase in HetR proteins or its activity (e.g. decreased level of PatS, a HetR inhibitor (43)) should be rejected based on the results that cyAbrB1 regulation of the genes is independent of HetR (Fig. 3A) and that expression of *hetZ*, a direct target of HetR (34), was not induced following *cyabrB1* knockdown (Fig 2). Upon nitrogen step-down, amounts of cyAbrB1 did not significantly decrease (Fig. S8). We propose that strong induction of HetR in proheterocysts (42) out-competes the repression of *hetP* and *hepA* by cyAbrB1 during heterocyst differentiation in a wild type strain. The view is partly supported by Figure 6, in which the over-expression of cyAbrB1 hampered the induction of *hepA* but not the induction of *hetP*. The observation could be explained by the fact that the binding affinity of HetR for the *hetP* promoter is much higher than that for the *hepA* promoter (38). Decrease of amounts of cyAbrB1 or cyAbrB1 activity attributed to glutathionylation (44) in proheterocysts is a potential reason. Taken together, we propose that a balance between the amounts of cyAbrB1 and HetR is a key determinant for the initiation of heterocyst differentiation.

It is typical that both a global activator and a repressor regulate prokaryotic cell differentiation. In *Streptomyces*, a global transcriptional activator AdpA and repressor BldD control morphological differentiation (45, 46), while in *Bacillus subtilis*, an activator Spo0A and a repressor AbrB regulate spore formation (47). Notably, both the DNA-binding domain of cyAbrB (AbrB-like family, Pfam14250) and that of AbrB from *B. subtilis* (MazE_antitoxin family, Pfam04014) belong to the same clan (AbrB, CL0132), although the former is located at the C-terminus and the latter is located at the N-terminus. However, the relationship between the activator and the repressor is different between *Anabaena* and *B. subtilis*, as well as *Streptomyces*. The expression of AdpA is regulated by BldD in *Streptomyces* (46, 48) and the expression of AbrB is regulated by Spo0A in *B. subtilis* (47). In contrast, our results demonstrated that HetR does not regulate cyAbrB1, and vice versa. Many cells differentiate in *B. subtilis* and *Streptomyces* simultaneously. Hence, the fact that one regulator governs the whole network is a practical strategy for cell differentiation in the above organisms. In contrast, only one-tenth of cells differentiate into heterocysts and the remaining vegetative cells maintain viability and photosynthetic activity in *Anabaena*. In addition, heterocyst differentiation is terminal (non-reversible) while cell differentiation in *B. subtilis* and *Streptomyces* is non-terminal (reversible). Therefore, robust inhibition of heterocyst differentiation by cyAbrB1, whose functioning is independent of HetR, could be an essential safety device in *Anabaena*, in concert with HetR inhibitors PatS (43) and HetN (49, 50) and regulation by HetR phosphorylation (51).

### Repression of heterocyst formation in ammonium medium

Heterocysts were formed in the presence of nitrate but not in the presence of ammonium in *cyabrB1* knockdown mutant C104 even though the expression of *hetP* was induced in the presence of ammonium similar to in the presence of nitrate when *cyabrB1* was knocked down. The results suggest the existence of an unidentified mechanism that regulates heterocyst differentiation. A previous study demonstrated that the over-expression of *hetP* induced heterocyst formation even in the presence of ammonium (33). The inconsistency between our results and those of the previous study could be explained by the unidentified mechanism, which would be cyAbrB1 dependent.

### Perspectives

In the present study, we could not identify target genes of cyAbrB1 other than *hetP* and *hepA,* which are not conserved in many non-heterocystous cyanobacteria. Up-regulation of the genes in a cyAbrB1 knockdown mutant could not explain why *cyabrB1* is essential in *Anabaena*. A transcriptome analysis would identify other target genes comprehensively, and would answer our question on whether cyAbrB1 conserved in cyanobacteria regulates core genes (10, 11). If cyAbrB1 regulated core genes, whether cyAbrB1 is involved in the reconstruction of metabolism during heterocyst development such as inactivation of photosystem II (52) or enhancement of photosystem I (53) would be an interesting question.

Our results demonstrated that cyAbrB1, but not cyAbrB2, specifically regulates the expression of *hetP* and *hepA*. However, heterocyst formation in the presence of nitrate was enhanced in double mutant C104B2 than in the single mutant C104 (Fig. 4). The results indicate that cyAbrB proteins can function both specifically and redundantly/cooperatively in *Anabaena*, as has been suggested in *Synechocystis* (20), although the underlying mechanism remains to be elucidated. A transcriptome analysis of C104B2, C104, and that of a *cyabrB1* knockdown mutant in *Synechocystis* could shed more light on the evolution and adaptation of cyanobacteria.

## Materials and Methods

### Bacterial strains and growth condition

*Anabaena* strains were cultured at 30°C under 30-35 μmol photons m^−2^ s^−1^ in BG11 medium (54) (17. 6 mM sodium nitrate as nitrogen source). BG11_0_ (lacking nitrogen sources) or BG11_a_ (5 mM ammonium chloride as nitrogen sources) were used after washing the cells twice with BG11_0_, where indicated. Each medium was supplemented with 20 mM HEPES-NaOH (pH 7.5). Two μg/ml each of spectinomycin and streptomycin and 25 μg/ml neomycin-sulfate were added when required. Liquid cultures were bubbled with air containing 1.0% (v/v) CO_2_ unless otherwise stated.

### Plasmid construction

Plasmids for the knockdown of *cyabrB1* by CRISPRi or the over-expression of *cyabrB1* were constructed using the hot fusion method (55). DNA fragments were inserted between the BamHI and KpnI sites of a genome-integrating vector pSU102-cyaA (26). Schematic representations of them are shown in Fig. S1 and detailed sequences are described in Supplemental information. Inactivation of *cyabrB2* was accomplished by replacing a 200-bp portion of the *cyabrB2* coding region with a spectinomycin resistant cassette as follows. Upstream and downstream regions of the *cyabrB2* gene were amplified by PCR using the primer pair 2080-5F and 2080-5R and the 2080-3F and 2080-3R pair, respectively. The spectinomycin cassette was inserted between upstream and downstream regions and the resultant construct was cloned between SacI and XhoI sites of pRL271 (56) to construct pR2080S. To construct the expression plasmids for the hexahistidine-tagged cyAbrB1 protein, DNA fragment containing the *cyabrB1* coding regions was amplified by PCR using the primer pair 0946-F and 0946-R. The amplified DNA fragment was cloned between NdeI and BamHI sites of the pET-28a expression vector (EMD Millipore) to construct pEcyAbrB1.

### RNA extraction and RT-qPCR analysis

Total RNA was extracted from cells using a phenol-based solution PGTX (57) and Zirconia/Silica beads (□ 0.1 mm). After treatment with DNase I (Takara Bio, Shiga, Japan), RNA was cleaned-up with NucleoSpin RNA kit (Takara Bio, Shiga, Japan). Synthesis of cDNA and qPCR were performed as described previously (26). Primers used in qPCR are listed in Table 1.

**Table 1.**
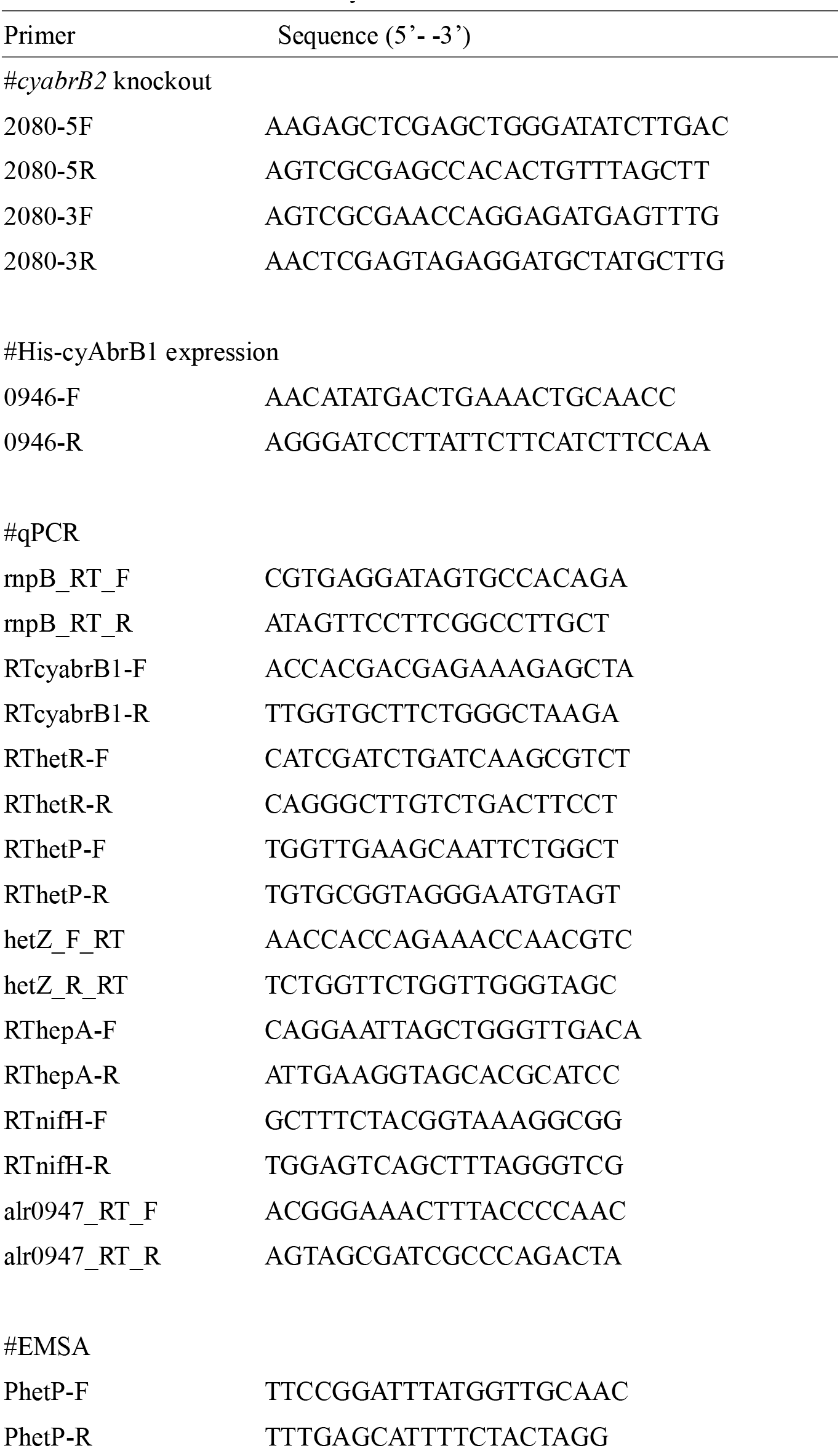
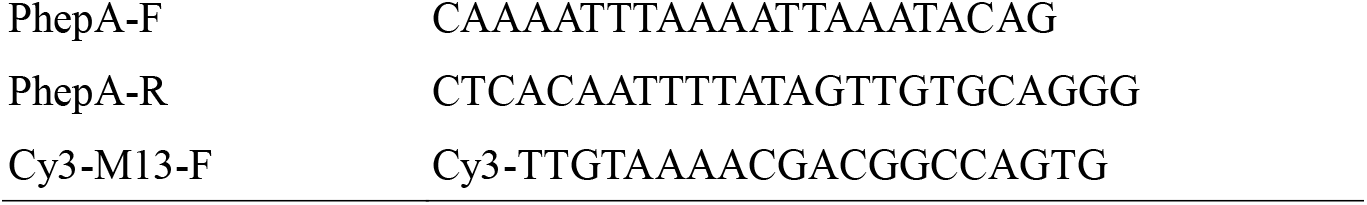
Primers used in this study

### Western-blotting analysis

Cell pellets were resuspended with 400 μl of extraction buffer [25 mM HEPES-NaOH (pH 7.5), 1 mM EDTA, 5 mM 2-mercaptoethanol and 1 × protease inhibitor cocktail (Roche)]. Cells were disrupted using Bullet Blender (Next Advance) in the presence of 0.5 g of stainless steel beads (□0.2 mm) with 3 cycles of agitation for 3 min and cooling for 3 min. Beads and cell debris were removed by centrifugation at 21,000 × *g* for 3 min. Western-blotting analysis was conducted as follows: Equal amounts of total protein were separated on a denaturing SDS-polyacrylamide gel and blotted onto a polyvinylidene fluoride (PVDF) membrane. cyAbrB1 was detected using a rabbit polyclonal antibody raised against His-cyAbrB1 from *Synechocystis* (20).

### Enrichment of heterocysts

The enrichment of heterocysts from *Anabaena* filaments was performed as described previously (26).

### Expression and purification of His-cyAbrB1

*E. coli* BL21(DE3) harboring pEcyabrB1 was grown at 37°C in 250 ml of Luria-Bertani medium. The recombinant gene was expressed in exponentially growing cells (an OD_600_ of 0.6) by adding 1 mM isopropyl-β-D-thiogalactopyranoside. After 5 h of incubation, the cells were harvested by centrifugation. His-cyAbrB1 was purified with the Ni-NTA Fast Start kit (Qiagen). The elution fractions containing the purified protein were loaded onto a PD MidiTrap G-25 column (GE healthcare) equilibrated with 20 mM phosphate buffer (pH 7.4) containing 0.5 M NaCl and 10% glycerol, and the protein was eluted with the same buffer.

### Gel mobility shift assay

The *hetP* promoter region was amplified by PCR using primer pair PhetP-F and PhetP-R and was cloned into the EcoRV site of the PBluescript II KS+ to construct pBPhetP. A Cy3-labeled probe PhetP was prepared by PCR using a Cy3-labeled M13-F primer and PhetP-R with pBPhetP as a template. His-cyAbrB1 was incubated with a Cy3-labeled probe (3 nM) in 20 μl of incubation buffer [20 mM Tris-HCl (pH 7.5), 10 mM MgCl_2_, 1 mM dithiothreitol, 40 ng/μl BSA, and 5% glycerol] for 30 min at room temperature. The mixtures were subjected to electrophoresis on a native 5% polyacrylamide gel and Cy3-labeled probes were detected on an FLA-9000 imaging system (FUJI Film). Non-labeled DNA probes PhetP, PhepA, and cyabrB1RT were prepared by PCR using primer pair PhetP-F and PhetP-R, primer pair PhepA-F and PhepA-R, and primer pair RTcyabrB1-F and RTcyabrB1-R, respectively.

## Supporting information

Sequence information

Suppemental Figure1

Suppemental Figure2

Suppemental Figure3

Suppemental Figure4

Suppemental Figure5

Suppemental Figure6

Suppemental Figure7

Suppemental Figure8

Suppemental Figure9

## Acknowledgments

This work was supported by the Institute for Fermentation, Osaka and by a Grant-in-aid for Scientific Research (C) 18K05395 from the Japan Society for the Promotion of Science.

---

Figure S1. Schematic representation of the genetic backgrounds of strains used in the present study. Components related to CRISPRi system or induction of *cyabrB1* were integrated at neutral site *cyaA* locus by single homologous recombination. Note that sizes of genes in the map are not proportional to sequence lengths.

Figure S2. Knockdown of *cyabrB1* leads to formation of heterocysts in the presence of nitrate, but not in the presence of ammonium. Cells were cultured for 48 h in the absence or presence of aTc and micro-photographed. Arrowheads indicate heterocysts. (A) *cyabrB1* knockdown strain (C104) was cultured in nitrate-containing medium and bubbled with air containing 1% (v/v) CO_2_. (B) *cyabrB1* knockdown strain (C104) was cultured in ammonium-containing medium bubbled with air containing 5% (v/v) CO_2_.

Figure S3. Knockdown of *cyabrB1* using sgRNA targeting different sites of *cyabrB1* led to formation of heterocysts in the presence of nitrate. Cells were cultured for 48 h in the absence or presence of aTc and micro-photographed. Arrowheads indicate heterocysts. Each *cyabrB1* knockdown strain was cultured in nitrate-containing medium bubbled with air containing 5% (v/v) CO_2_.

Figure S4. Confirmation of *cyabrB1* knockdown. (A and C) After 48 h of cultivation in the absence or presence of 50 ng/ml aTc, RNA was extracted and RT-qPCR was performed. *rnpB* was used for normalization. Data represent mean ± SD (n = 3 from independent culture). Amounts of each gene relative to that in C100 grown in nitrate-containing medium bubbled with air containing 5% (v/v) CO_2_ in the absence of the inducer are shown. (B and D) After 48 h of cultivation in the absence or presence of 50 ng/ml aTc, total protein was extracted and western blotting using anti-cyAbrB1 antibody was performed. (A and B) Each strain was grown in nitrate-containing medium and bubbled with air containing 5% (v/v) CO_2_. (C and D) Strain C104 was cultured in nitrate-containing medium bubbled with air containing 5 or 1% (v/v) CO_2_ or in ammonium-containing medium bubbled with air containing 5% (v/v) CO_2_.

Figure S5. Expression of *nifH* encoding nitrogenase subunit in *cyabrB1* knockdown strain C104 under different conditions. After 48 h of cultivation under the indicated conditions in the absence or presence of 50 ng/ml aTc, RNA was extracted and RT-qPCR was performed. *rnpB* was used for normalization. Data represent mean ± SD (n = 3 from independent culture). Amounts of *nifH* relative to that in C100 grown in nitrate-containing medium bubbled with air containing 5% (v/v) CO_2_ in the absence of the inducer are shown.

Figure S6. Knockdown of *cyabrB1* in *cyabrB2* deletion background. Cells were cultured in nitrogen-containing media bubbled with air containing 1% (v/v) CO_2_ in the absence or presence of aTc for 48 h. (A) Confirmation of *cyabrB1* knockdown. Cells were cultured in the presence of nitrate. RNA was extracted and RT-qPCR was performed. *rnpB* was used for normalization. Data represent mean ± SD (n = 3 from independent culture). Amounts of *cyabrB1* relative to that in C100 grown in nitrate-containing medium bubbled with air containing 5% (v/v) CO_2_ in the absence of the inducer are shown. (B) Confirmation of cyAbrB1 knockdown. Each strain was grown in the presence of nitrate or ammonium-containing medium. Total protein was extracted and western blotting using anti-cyAbrB1 antibody was performed. (C) No heterocyst formation of 104B2 following *cyabrB1* knockdown in ammonium-medium. Cells of C104B2 were cultured in the presence of ammonium and were micro-photographed.

Figure S7. Heterocyst formation in *cyabrB1* knockdown strain C104 in the absence of combined nitrogen sources. Nitrate was depleted from the culture medium in strain C104. Subsequently, C104 was cultured in nitrogen-free medium in the absence or presence of 50 ng/ml aTc. Images were photographed after 24 (A) or 48 h (B) of cultivation. Arrowheads indicate heterocysts. (C) Lengths of vegetative cell intervals between heterocysts in C104 cultured in nitrogen-free medium for 48 h in the absence (gray bars) or presence of aTc (black bars). (D) Confirmation of cyAbrB1 knockdown under nitrogen-fixation conditions. After 48 h of cultivation in nitrogen-free medium in the absence or presence of 50 ng/ml aTc, total protein was extracted and western blotting using anti-cyAbrB1 antibody was performed.

Figure S8. Expression of cyAbrB1 in wild type background. (A and B) Expression of cyAbrB1 after nitrogen depletion. Cells of PCC 7120 cultured in nitrate medium were washed two times with nitrogen-free medium. Subsequently, cells were resuspended in nitrate-medium (+N) or nitrogen-free medium (−N) and were cultured for 8 h. (A) Total protein was extracted and western blotting using anti-cyAbrB1 was performed. (B) Relative amounts of cyAbrB1 compared to that before nitrogen depletion were quantified from western blots. Data represent mean ± SD (n = 3 from independent culture). (C and D) Expression of cyAbrB1 in heterocysts. (C) After 24 h of nitrogen depletion, heterocysts were enriched. Total protein was extracted from whole filaments (lane W) and enriched heterocysts (lane H). Western blotting using anti-cyAbrB1, anti-RbcL, and anti-NifH antibodies was performed. RbcL and NifH are marker proteins for vegetative cells and heterocysts, respectively. (D) Relative amounts of cyAbrB1 in heterocysts compared to that in whole filaments were quantified from western blots. Data represent mean ± SD (n = 3 from independent culture).

Figure S9. Effect of *cyabrB1* knockdown or over-expression on expression of *alr0947*. RNA was extracted and RT-qPCR was performed. *rnpB* was used for normalization. Data represent mean ± SD (n = 3 from independent culture). (A and B) Cells were grown in the absence or presence of aTc for 48 h. (A) Each strain was grown in nitrate-containing medium bubbled with air containing 5% (v/v) CO_2_. (B) C104 was grown under indicated conditions. Amounts of *alr0947* relative to that in C100 cultured in nitrate-containing medium bubbled with air containing 5% (v/v) CO_2_ in the absence of the inducer are shown. (C) Each strain was grown in nitrogen-free medium for 8 h. Data represent mean ± SD (n = 3 from independent culture). Amounts of *alr0947* relative to that in C100 cultured in nitrate-free medium in the absence of the inducer are shown.

