## Supplementary material for "cyAbrB transcriptional regulators as safety devices to inhibit heterocyst differentiation in *Anabaena*": Sequence information

```
Supporting information
Detailed sequences of promoters and genes used in this study.
#sgRNA (-35 and –10 regions: underlined, guide sequence: italicized)
```

>*P_J23119_-sgRNA-1* (for C104)

TTGACAGCTAGCTCAGTCCTAGGTATAATGCTAGC

*TGCTTTTCCAGTTAAAGGCG*

GTTTTAGAGCTAGAAATAGCAAGTTAAAATAAGGCTAGTCCGTTATCAAC

TTGAAAAAGTGGCACCGAGTCGGTGCTTTTTT

>*P_J23119_-sgRNA-2* (for C105)

TTGACAGCTAGCTCAGTCCTAGGTATAATGCTAGC

*TGTTTAGCTCTTTCTCGTCG*

GTTTTAGAGCTAGAAATAGCAAGTTAAAATAAGGCTAGTCCGTTATCAAC

TTGAAAAAGTGGCACCGAGTCGGTGCTTTTTT

>*P_J23119_-sgRNA-3* (for C106)

```
TTGACAGCTAGCTCAGTCCTAGGTATAATGCTAGC
TAGCATCTGCATAAAGTTAC
GTTTTAGAGCTAGAAATAGCAAGTTAAAATAAGGCTAGTCCGTTATCAAC
TTGAAAAAGTGGCACCGAGTCGGTGCTTTTTT


#cyabrB1 (-35 and –10 regions, transcriptional start site: underlined, coding region of cyabrB1: green)
```

>*P_L03_-cyabrB1*

TCCCTATCAGTGATAGAGATTGACATCCCTATCAGTGATAGATATAATGGCCACATCAG

AATTCATCAGTTTGGAGAGAGTAAGGAAATTTTGAGCGTA

ATGACTGAAACTGCAACCGCGCCTTTAACTGGAAAAGCACTGCTAGCTAAGGTAAAAGAACTTTCTACTCTACCACGACGAGAAAGAGCTAAACAGTGCGGTTACTATACCGTTACTAAGAATAACCAAGTCCGGGTTAATCTAACAGATTTCTATGATGCTTTGCTATCGGCTAGAGGTATTCCTCTTAGCCCAGAAGCACCAAAAGATGGTCGTGGTCGTGAACCGACATACCGAGTTAGCGTTCATCAAAATGGTCAAATTGTGATTGGTGCTACTTACACCAAAGCAATGGGACTAAAACCAGGTGATGAATTTGAAATTAGATTGGGATACAAGCATATTCACTTGATTCAACTCGGTGAAAGCGATAAAAAGCTCTCCCCAGATTTAGATATCGACGAATCTGATGAAGATTTGGAAGATGAAGAATAA

ATTCTCTGAGGAAGGGATAAGTTAAAGTGATTACTTTTAACGTGTCCCTACTACCATATCAATCTGGTATCAGAGATTAAATAATTTTATCTAACGCAAAAACTACGATTTATTCCTTGCAGAAGTGACTACTCTTAGTAGAGGCTTGATTTATAAGCCAAAAATTTATTCTGCAAATAGTCTACCGTGATTTTTGTGTT
