## Supplementary figures and images for "cyAbrB transcriptional regulators as safety devices to inhibit heterocyst differentiation in *Anabaena*"

### Suppemental Figure1

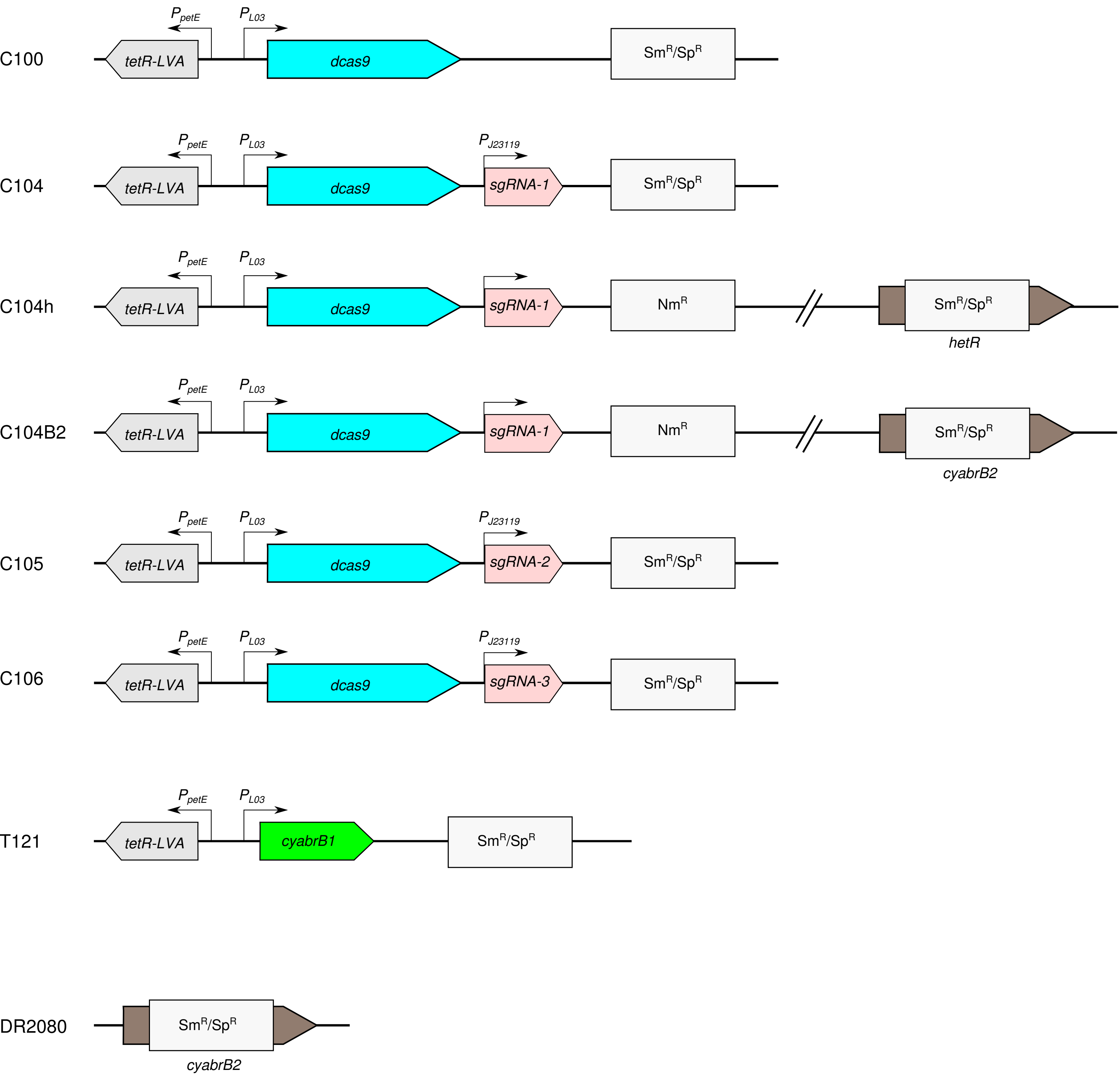

### Suppemental Figure2

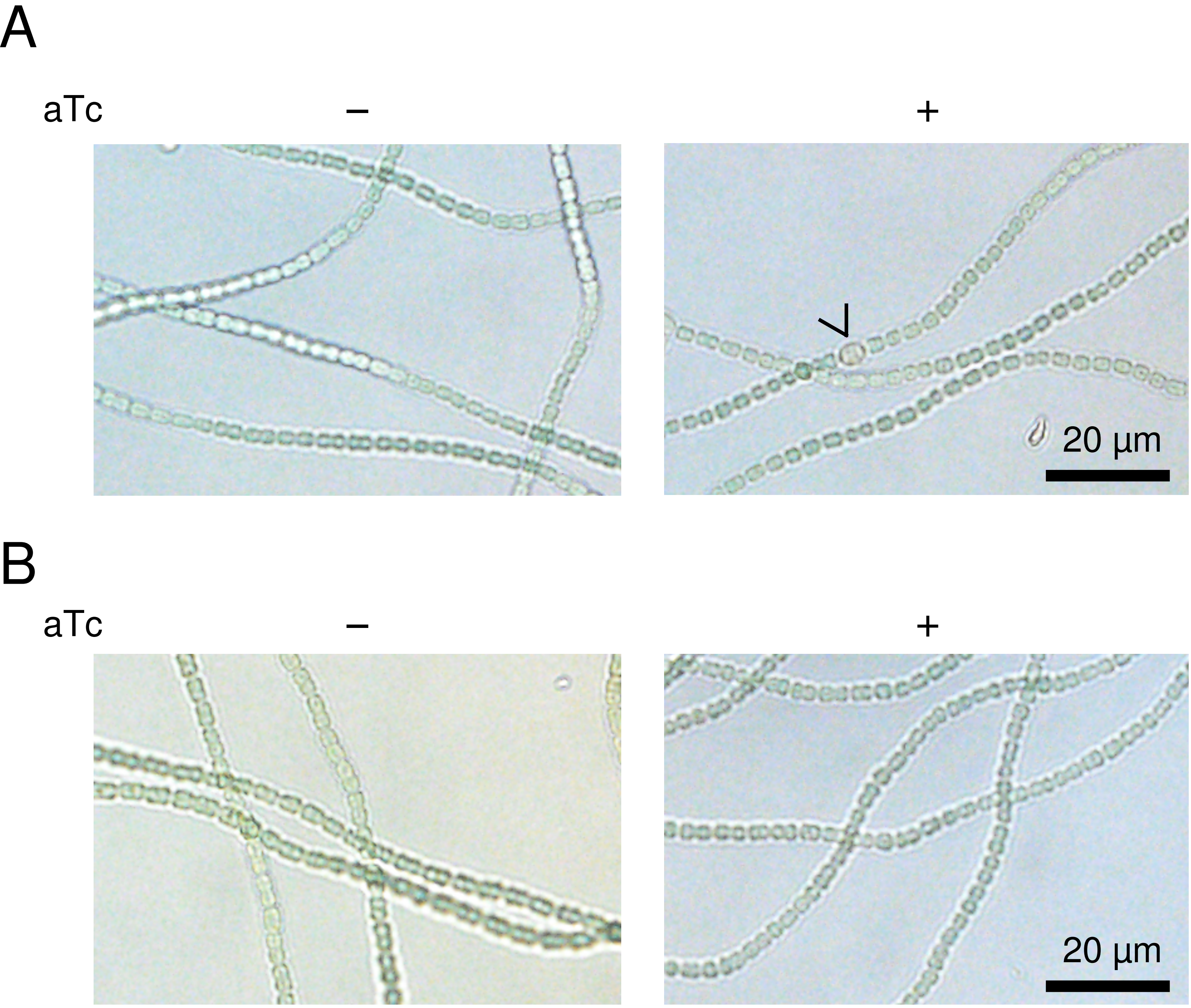

### Suppemental Figure3

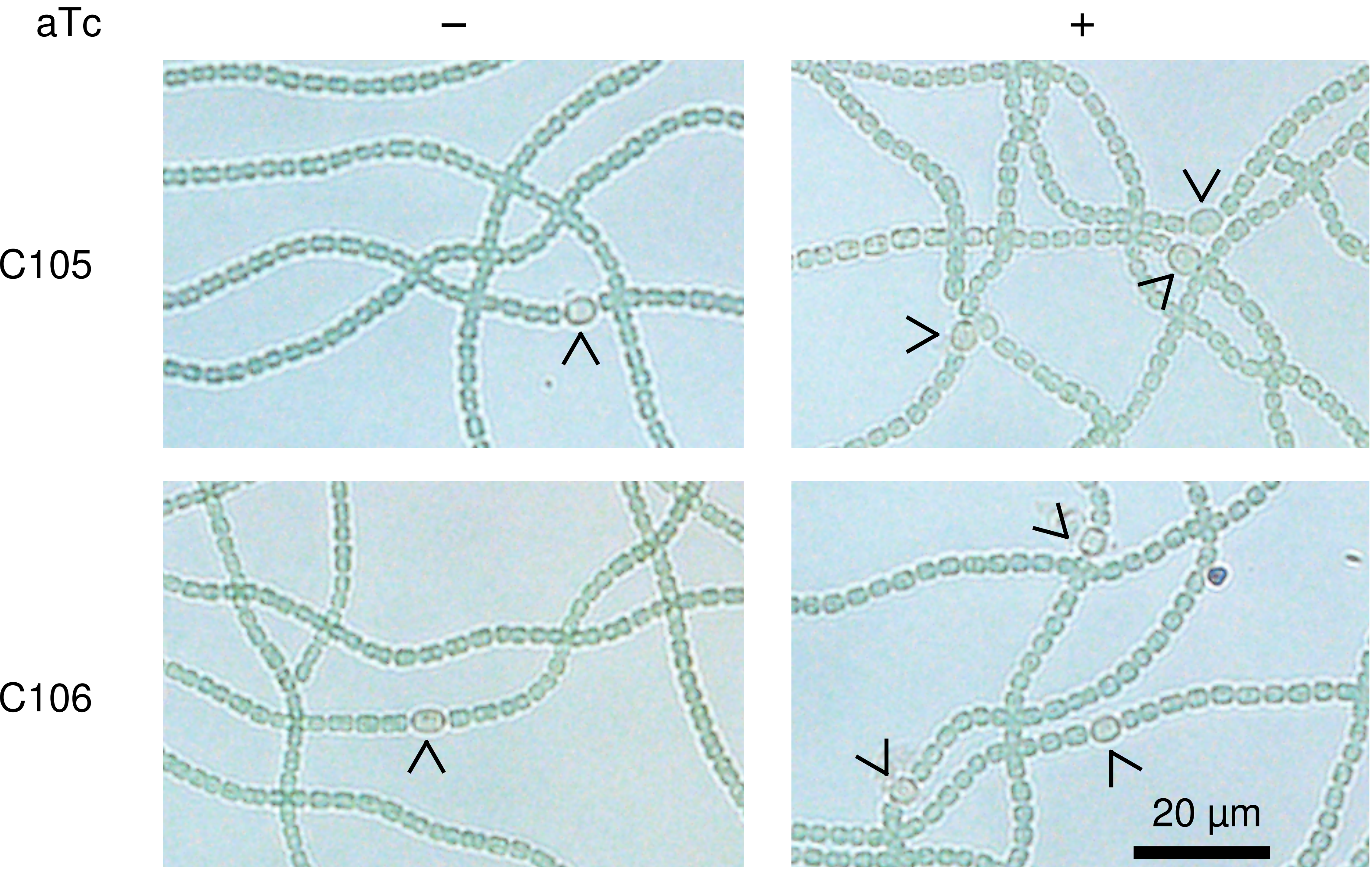

### Suppemental Figure4

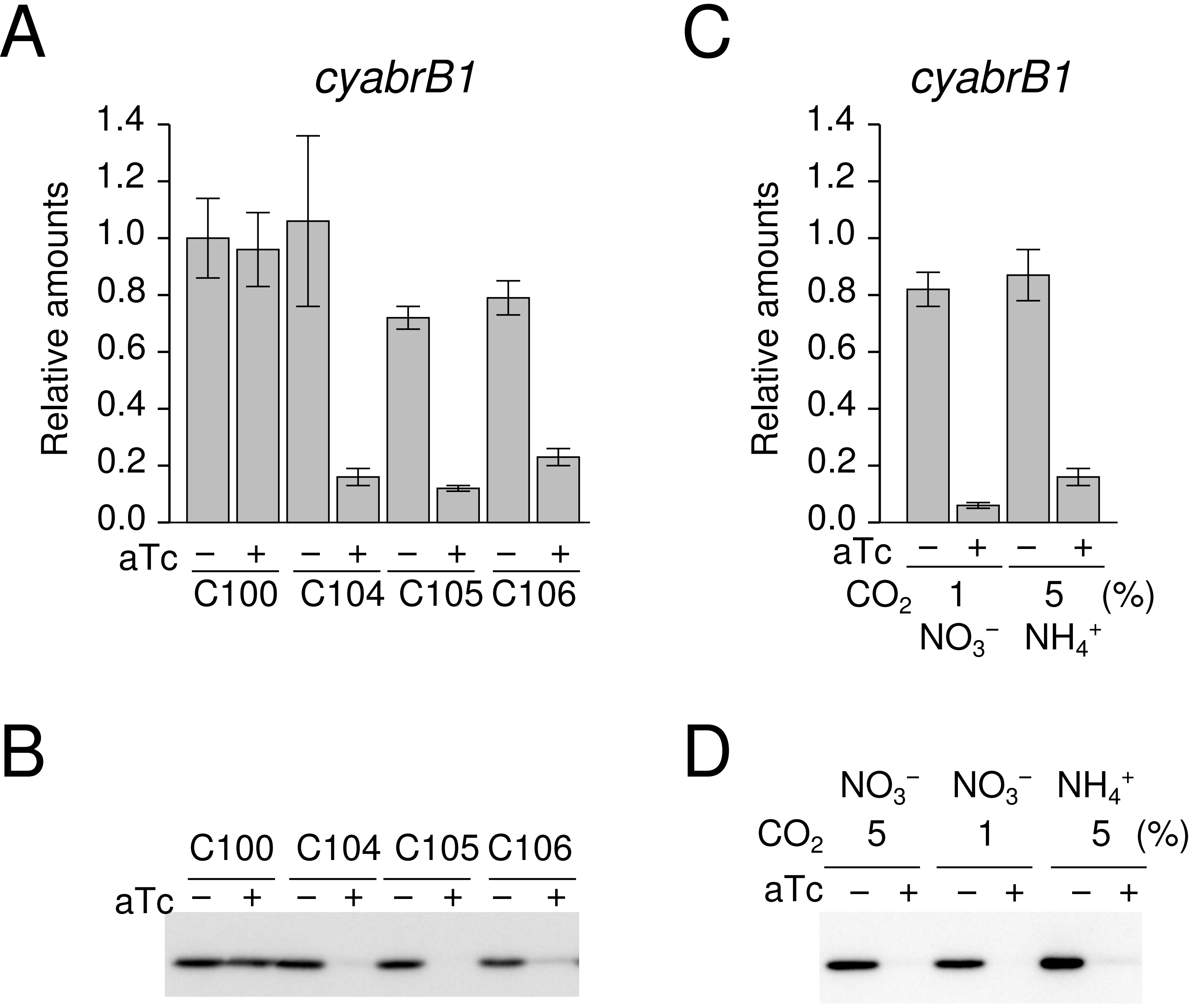

### Suppemental Figure5

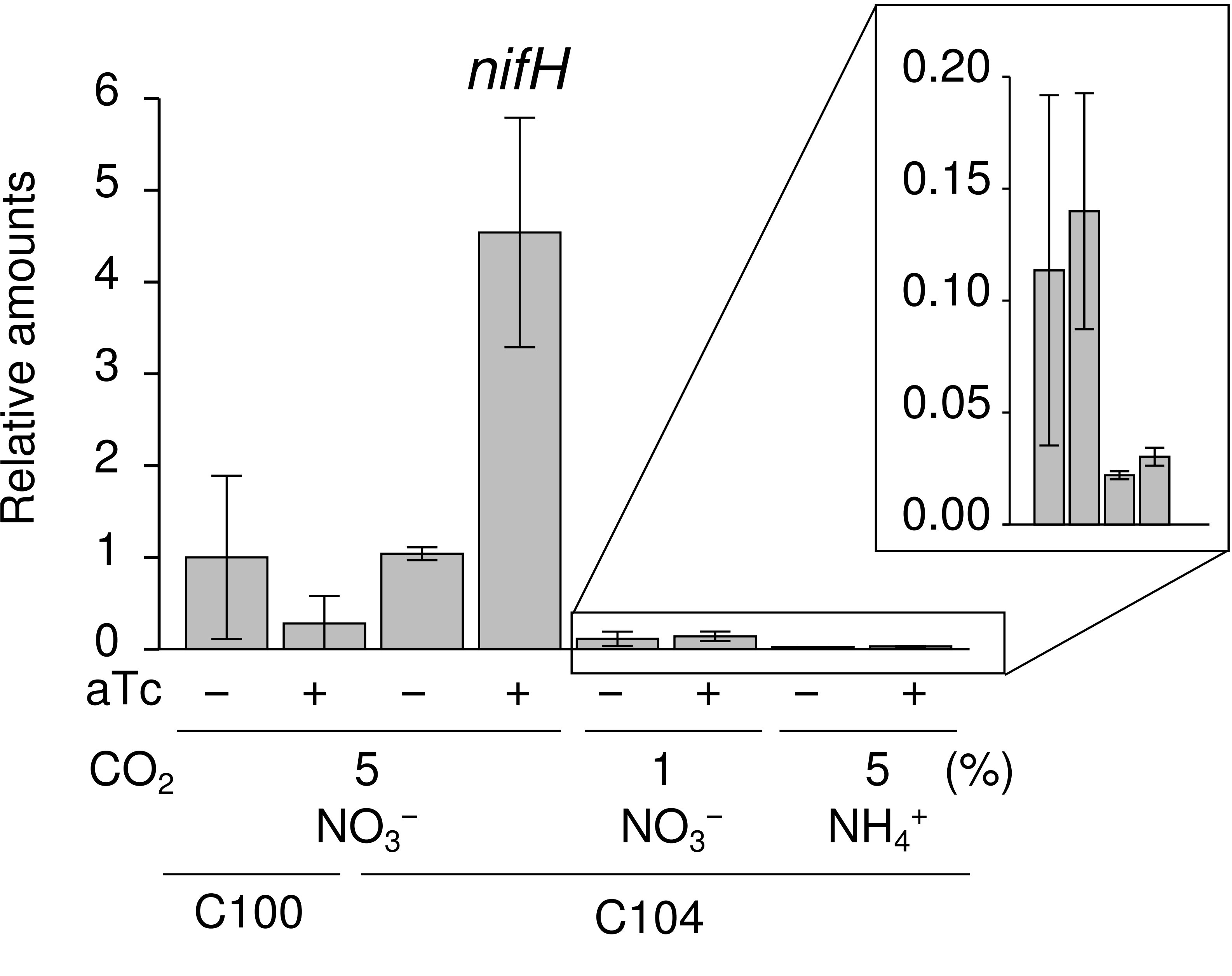

### Suppemental Figure6

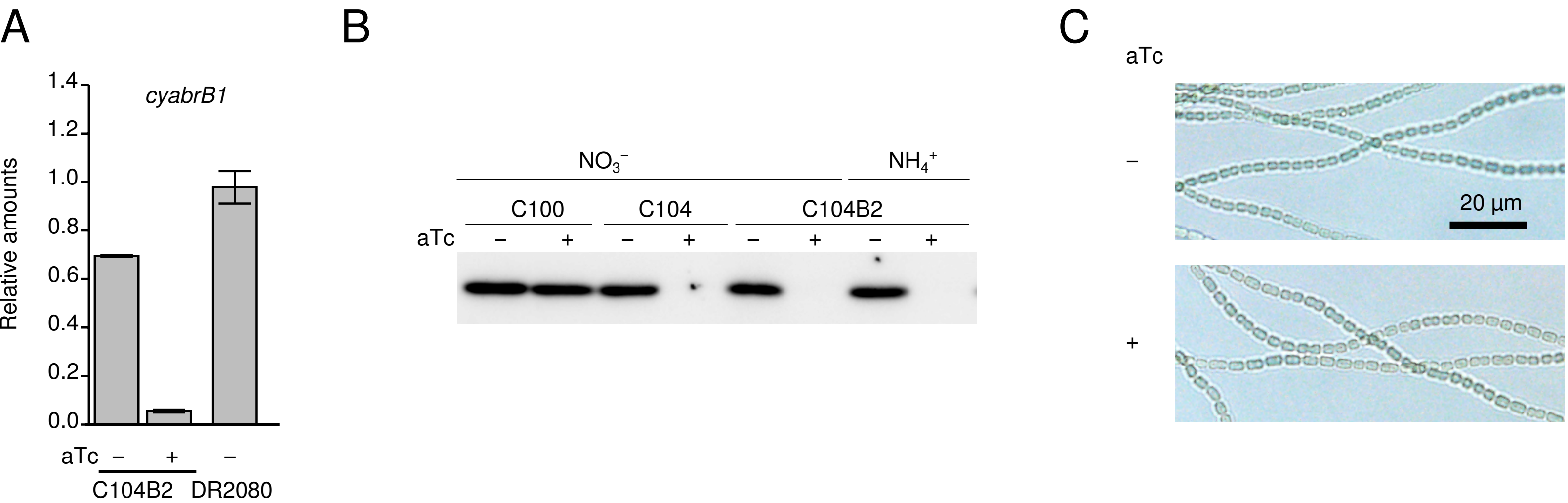

### Suppemental Figure7

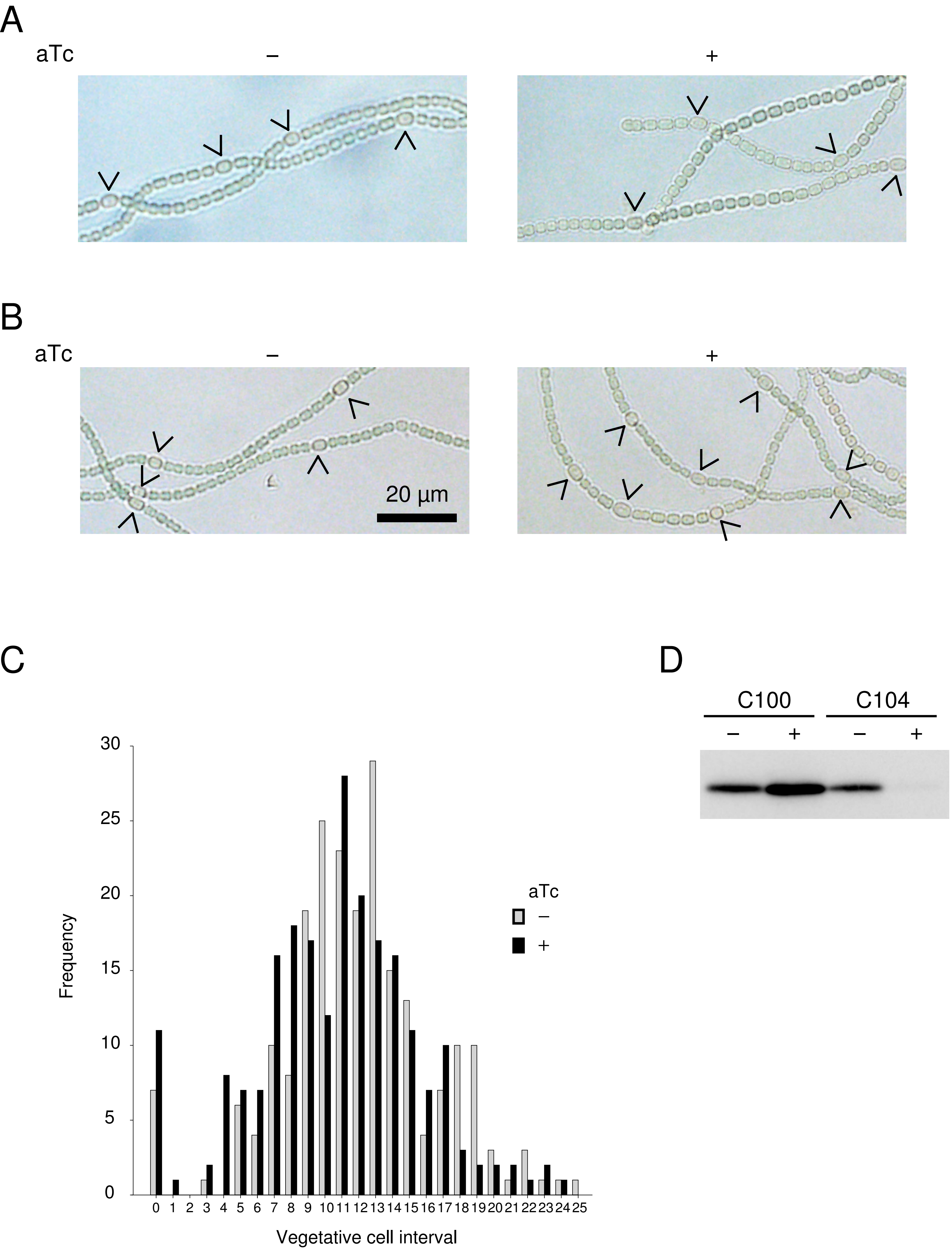

### Suppemental Figure8

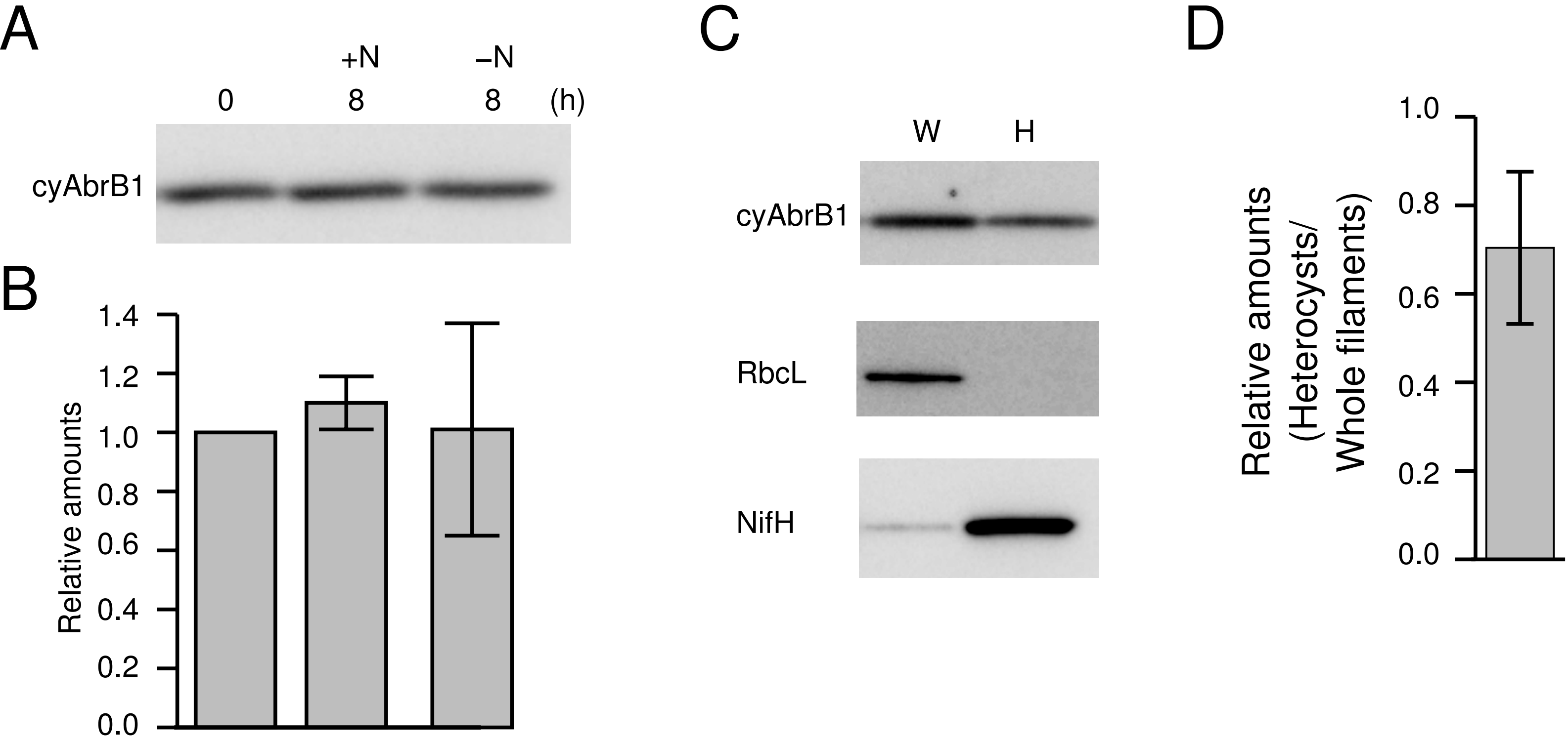

### Suppemental Figure9

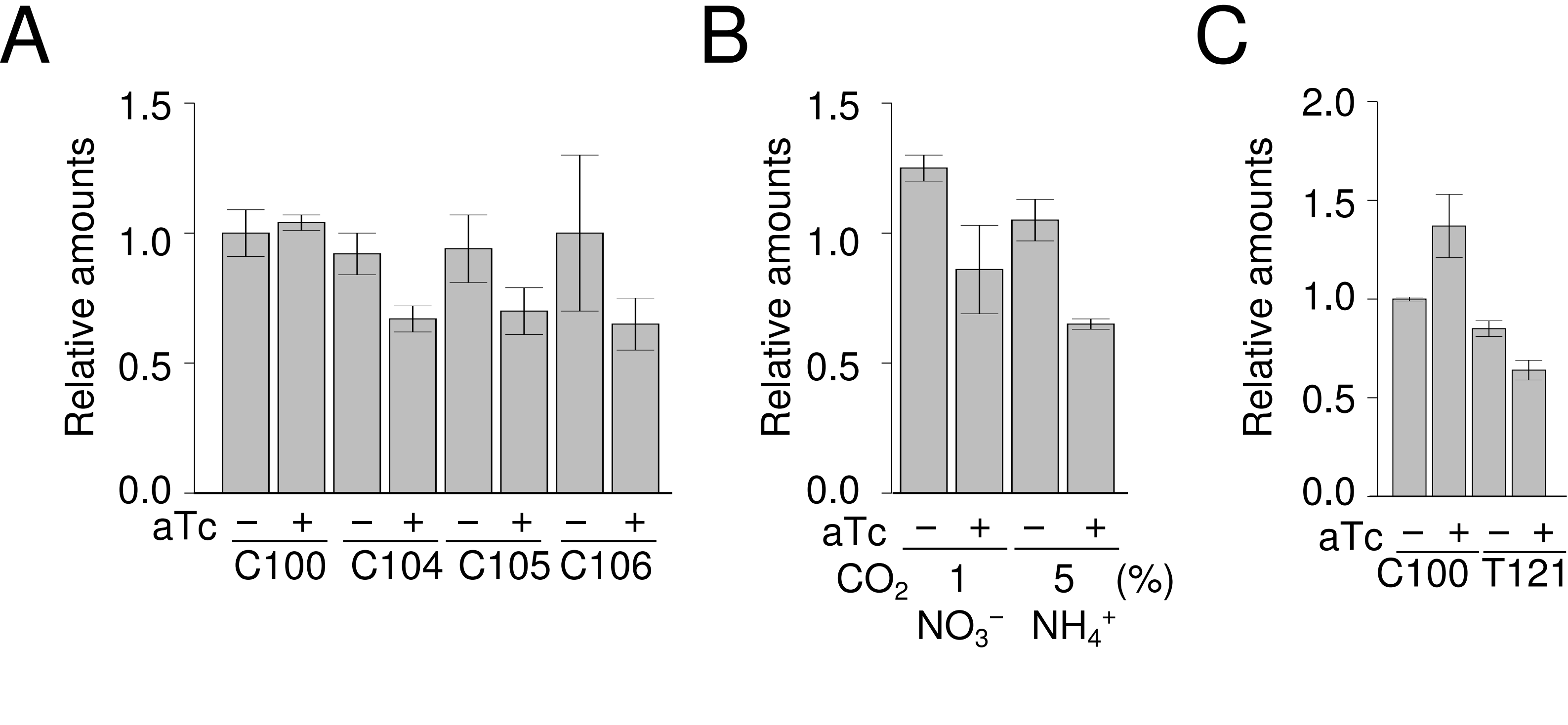
